# The Asian tiger mosquito, *Aedes albopictus* (Skuse, 1894), a vector of dengue, chikungunya and zika, reaches Portugal

**DOI:** 10.1101/192575

**Authors:** Eduardo Marabuto, Maria Teresa Rebelo

## Abstract

The mosquito *Aedes albopictus* is here reported for the first time in Portugal, from the south of the country, at least 240km west of the nearest known observation in Spain. A population of more than fifty specimens was spotted within a suburban garden over seven days of survey. As an important vector of Human affecting zoonoses such as dengue, chikungunya and yellow-fever, the presence of this mosquito in Portugal now enhances the outbreak chances for such diseases.

## Background

The Asian tiger mosquito, *Aedes albopictus* (Skuse, 1894) is a tropical species originally from south-eastern Asia [1]. It has experienced a rampant human-mediated range expansion since the 1970’s to now comprehend almost the whole of the tropical and subtropical areas of the world. As an eclectic haematophagous species, it attacks humans and is able to use a number of man-made and natural structures where stagnant water is present [2]. Eggs are able to survive for extended periods of time in complete dryness and diapause over unsuitable, cold season, making it an especially resilient species [1].

The worldwide spread of *A. albopictus* began with the Pacific and Indian islands, followed by North America in 1985, Brazil and other south American nations in subsequent years. Africa was colonised in 2000, despite isolated records since at least 1990 [3]. The closely related *Aedes aegypti* (Linnaeus, 1762) is a long known species in Europe, reported in mainland Portugal until the 1950’s, and now established in the island of Madeira, where it has been responsible for heightened public-health issues [4,5]. *A. albopictus* only reached Europe in 1979, arriving to Albania presumably as eggs or larvae in used, water-filled tyres [6]. It has now expanded through the Peri-Mediterranean area [7], east to Georgia [8] and west to the remaining of southern Europe.

After fully colonising Italy and the Mediterranean coast of France, the earliest report from the Iberian Peninsula is from 2004, when the species was found in Catalonia [9]. Here, the spread of the species progressed particularly fast along the warmer, Mediterranean shoreline. The latest colony of *A. albopictus* to be reported was found in 2015, near Algeciras in central Andalucía province [10]. This is about 240 km away from the Portuguese border along the Atlantic coast. The presence of the mosquito itself had long been antecipated in Portugal [3], warned for in the national news during 2016 but the mosquito had eluded all the intensive surveying efforts by both Spanish scientists working along the Huelva coast [10] and the Portuguese REVIVE (National Network for Vector Surveillance) team throughout the country [11].

## First contact

The first observation of this species in Portugal corresponds to several females foraging on human blood on the 31-VII-2017 in the late afternoon in a private condominium near the golf resort of Vila Sol, Vilamoura, Faro, Algarve, Portugal (coordinates: 37.090, −08.093) (Fig. 1).

**Figure 1.**
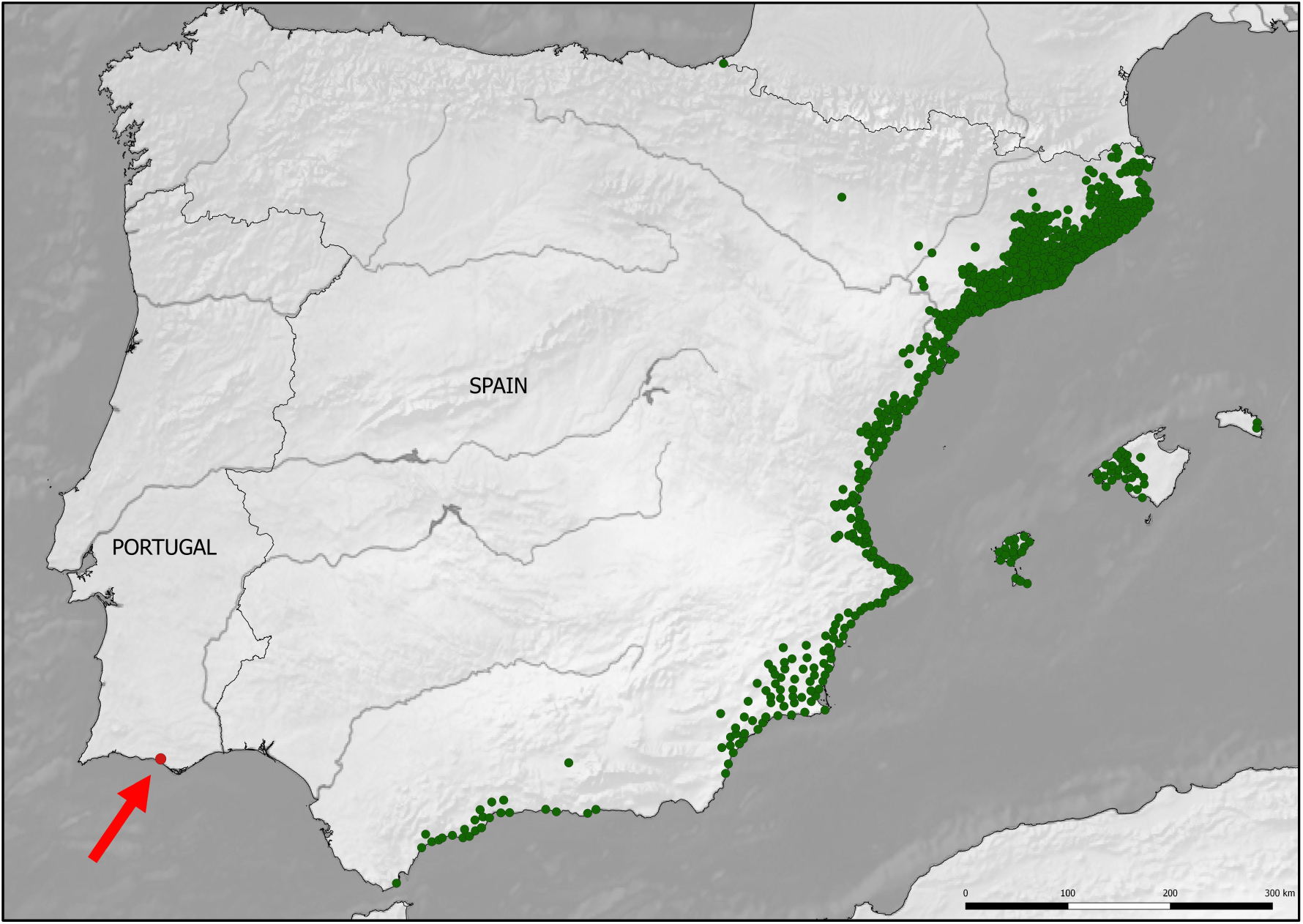
Distribution of *Aedes albopictus* in the Iberian Península. In green, distribution in Spain according to Collantes et al. (2016). In red, highlighted by arrow, new observation in Portugal.

The site is an anthropogenically engineered subtropical garden with a prevalence of alochthonous flora surrounded by villas and including several swimming-pools. At dusk, several garden lights at ground level are regularly lit and during the night, an automatic irrigation system maintains humidity levels high, even during the summer. These favourable conditions, lacking local ecological elements such as predators favour the presence of alien species, like the African mantis *Miomantis paykullii*, [12]. Not uncommonly, artificial structures are filled with water for long periods of time and the golf-course nearby holds several permanent artificial fresh-water bodies.

## Further observations

From 31-VII to 6-VIII-2017, 58 *Aedes albopictus* adult specimens, both males and females, were observed or collected with a tube, either foraging on human blood (Fig. 2) or resting among the vegetation. One female was seen ovipositing in the nearby swimming-pool margin, despite its regular maintenance.

**Figure 2.**
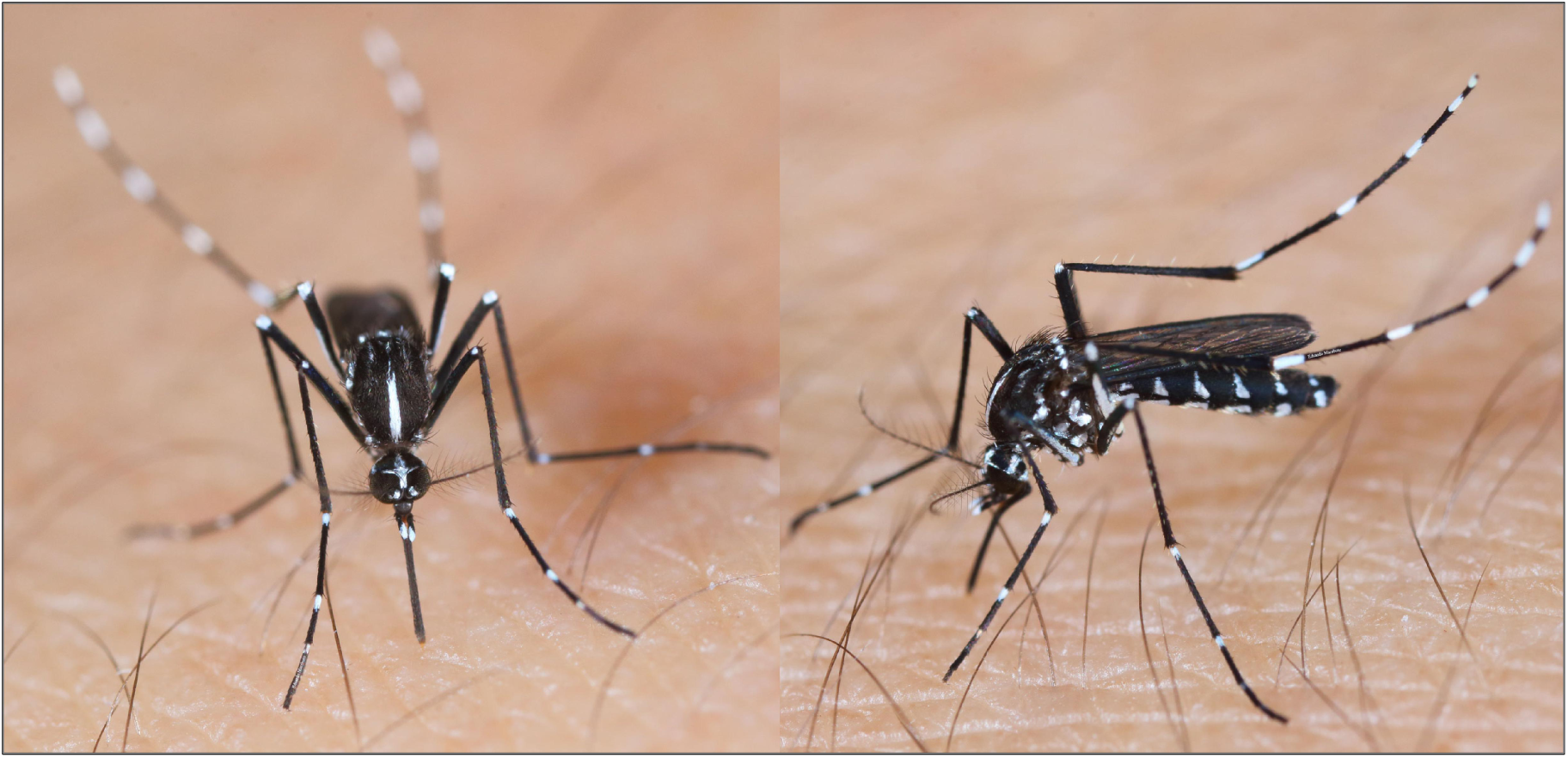
*Aedes albopictus* female specimen in Portugal, foraging on human blood. Note characteristic pronotal white stripe, distinct silver white scales on the palpi and tarsi, as well as narrow scales over wing root and basal abdominal tergal markings not connected with lateral ones.

Observation of Asian tiger mosquitos *in situ* revealed an agressive behaviour towards human subjects, in line to observations by Halasa *et al.* [13]. Specimens arriving and settling for feeding on human blood, always in the same place, for a period of 3 days revealed a peak of activity in the late afternoon, which ceased just before complete obscurity (21:30h), and a high population density (Fig. 3).

**Figure 3.**
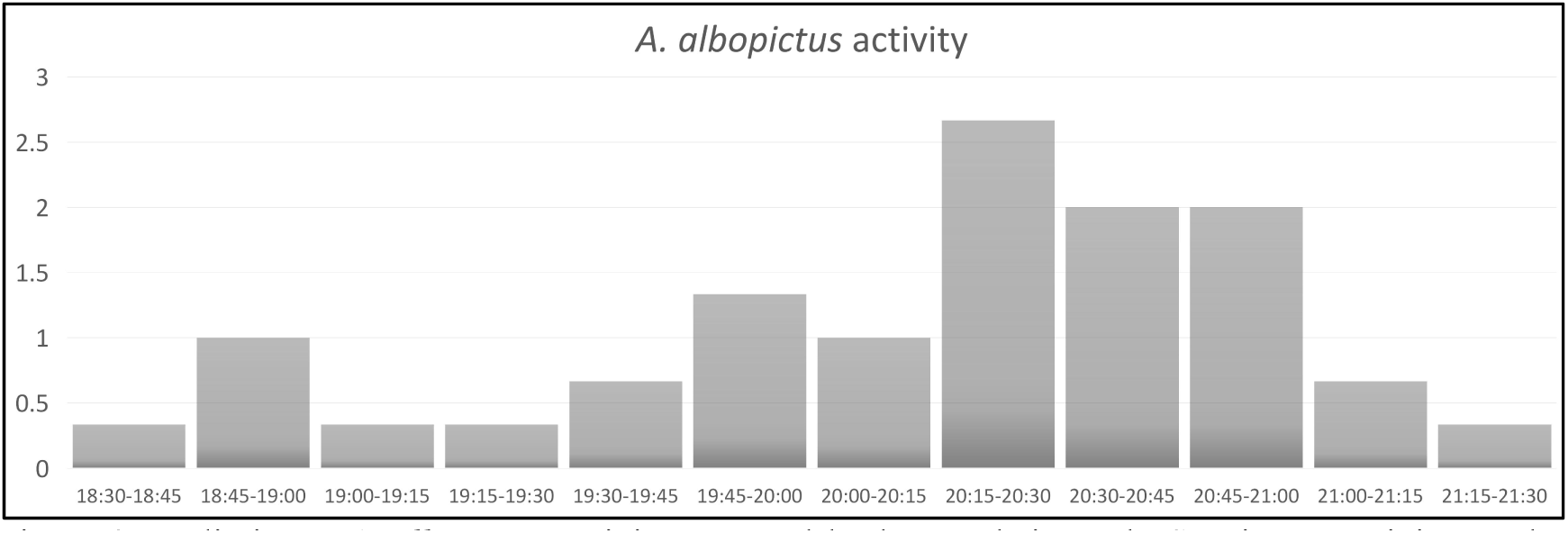
Preliminary *A. albopictus* activity assessed by human bait catch. Specimens arriving at the bait’s vicinity were captured to avoid recapture, every 15 minutes. X axis is a timeline of 15’ interval, y axis is average of the 3 days of sampling.

## Conclusions

*Aedes albopictus* is an ecologically aggressive and plastic species, evidently able to cope with and benefiting from Human interference upon natural ecosystems [14]. In Europe since 1979, it has progressively colonised suitable, usually anthropogenic habitats, from Georgia to the Iberian Peninsula.

With its presence confirmed in Portugal, this mosquito’s journey has reached the Atlantic shores and one of the most suitable areas for the species [7,15]. This suitability is further enhanced by the proliferation of man-made subtropical-type gardens and predator-free water bodies, available year-round.

The ecological and social disturbance caused by this species would not be very different from the many others currently arriving and establishing within the Mediterranean area, often with ecological impact, and it would pass relatively unnoticed if *A. albopictus* was not a well-known vector of several arboviruses.

In fact, throughout the tropical areas of the world and occasionally outside these regions, following travel of infected hosts, this species is associated with the transmission of more than 20 arboviruses, among which is dengue, yellow fever, chikungunya and even zika [16, 17, 18].

Outbreaks of these zoonoses have already taken place in the newly invaded range of *Aedes albopictus* and *A. aegypti*, with these species involved as vectors. Major nearby events such as the dengue outbreak in Madeira [5] and of chikungunya in Italy [19] must be taken into account for they are now possible in mainland Portugal. Despite the intense monitoring over this expanding vector and knowledge of the viruses it may transmit, the reported increasing risk of autochthonous transmission of dengue, chikungunya and zika viruses by Spanish health authorities [20] is very real to Portugal as well.

Strategies for prevention and control of this epidemiologically aggressive invader, emphasizing Integrated Vector Management as a multi-sectorial approach, should reinforce linkages between health and environment fields, providing stakeholders with a complete tool for improving decision capacity. Combined assessment of potential introduction gateways and local climatic zones may form an evidence base for planning efficient mitigation strategies against *Aedes albopictus*.

## Acknowledgements

We would like to thank all involved family members of the first author upon finding of the in discussing aspects related to the occurence of the species, their location and aprehension during the time-frame of the study, especially Manuel Marabuto for allowing their photographic capture while foraging. First author’s research is supported by FCT (Fundação para a Ciência e Tecnologia), Portugal doctoral grant SFRH/BD/102801/2014.

